# Loss of genital epithelial barrier function is greater with depot-medroxyprogesterone acetate than intravaginal rings that release etonogestrel and ethinyl estradiol

**DOI:** 10.1101/2023.01.25.525538

**Authors:** Mohan Liu, Rodolfo D. Vicetti Miguel, Nirk E. Quispe Calla, Kristen M. Aceves, Linda Fritts, Christopher J. Miller, John A. Moss, Marc M. Baum, Thomas L. Cherpes

## Abstract

The injectable progestin depot-medroxyprogesterone acetate (DMPA) is a popular contraceptive choice in sub-Saharan Africa although mouse models indicate it weakens genital epithelial integrity and barrier function and increases susceptibility to genital infection. The intravaginal ring NuvaRing® is another contraceptive option that like DMPA suppresses hypothalamic pituitary ovarian (HPO) axis function with local release of progestin (etonogestrel) and estrogen (ethinyl estradiol). As we previously reported that treating mice with DMPA and estrogen averts the loss of genital epithelial integrity and barrier function induced by DMPA alone, in the current investigation we compared genital levels of the cell-cell adhesion molecule desmoglein-1 (DSG1) and genital epithelial permeability in rhesus macaques (RM) treated with DMPA or a NuvaRing®re-sized for RM (N-IVR). While these studies demonstrated comparable inhibition of the HPO axis with DMPA or N-IVR, DMPA induced significantly lower genital DSG1 levels and greater tissue permeability to intravaginally administered low molecular mass molecules. By identifying greater compromise of genital epithelial integrity and barrier function in RM administered DMPA vs. N-IVR, our results add to the growing body of evidence that indicate DMPA weakens a fundamental mechanism of anti-pathogen host defense in the female genital tract.

## INTRODUCTION

Most women in sub-Saharan Africa (SSA) become pregnant before 18 years of age [1], an observation that indicates adolescent girls and young women (AGYW) in this region benefit from access to effective contraception. However, other epidemiological data from SSA shows that nearly 80% of the individuals between 10-19 years of age that acquire human immunodeficiency virus (HIV) infection each year are female [2]. These data propelled efforts to define the behavioral and biological variables that increase HIV susceptibility in AGYW, and results from several observational studies suggested risk of acquiring HIV is about 40% higher in women using the injectable progestin depot-medroxyprogesterone acetate versus women using no hormonal contraception [3-5]. These findings are especially relevant to SSA, where DMPA is utilized by nearly half of all contraceptors [6]. However, the validity of these results is questioned as no observational study had been specifically designed to define impact of DMPA on HIV susceptibility and study conclusions were potentially confounded by the higher frequency of unprotected sex among women using DMPA [7]. On the other hand, the Evidence for Contraceptive Options and HIV Outcomes (ECHO) Trial, a randomized trial that compared effects of 3 different contraceptives on HIV transmission, found that women using DMPA were at a 29% greater risk of HIV acquisition than women using a levonorgestrel (LNG)-releasing subdermal implant (P= 0.06) [8].

While clinical research continues to explore the association between HIV and DMPA, there is additional need to define the biological mechanisms that potentially underlie the connection between DMPA and HIV susceptibility. The genital epithelial barrier, a first-line defense that impedes virus entry at mucosal surfaces, is maintained by several cell-cell adhesion molecules, including the desmosomal cadherins desmoglein-1 (DSG1) and desmocollin-1 (DSC1), and we previously reported that DMPA comparably reduces genital DSG1 and DSC1 levels in mice and humans and this loss of epithelial integrity increases tissue permeability to penetration by low molecular mass (LMM) molecules [9, 10]. These results revealed mice accurately model genital epithelial changes that occur in women using DMPA and provided new biological plausibility for the putative connection between DMPA and HIV susceptibility. Subsequent clinical studies confirmed these results, including reports of loss of genital epithelial integrity and lower levels of DSG1 protein in ectocervical biopsy specimens from women using DMPA vs. no hormonal contraception [11, 12]. In further exploration of murine models, we also showed that DMPA respectively increases susceptibility of wildtype and humanized mouse to genital infection with herpes simplex virus type 2 (HSV-2) and cell-associated HIV-1, while treatment of mice with both DMPA and estrogen (E) preserves genital epithelial barrier function and confers resistant to viral infection [8, 13]. Based on these results, we posited that intravaginal rings (IVR) releasing etonogestrel (ENG) and ethinyl estradiol (EE) (e.g., NuvaRing®) are less likely than DMPA to impair genital epithelial integrity and barrier function. We tested this hypothesis in the current investigation by comparing genital DSG1 protein levels and genital permeability to LMM molecules in rhesus macaques (RM) administered DMPA or NuvaRings® that had been re-sized for use in RM (hereafter these modified IVR are identified as N-IVR).

## MATERIAL AND METHODS

### IVR modification

NuvaRings® (Organon, Jersey City, NJ) were purchased and modified to fit RM by removing a 66 mm length. Thermal welding was used to rejoin ends and form rings approximately 25 mm in diameter. While commercially available NuvaRings® contain 11.7 mg of ENG and 2.7 mg of EE, N-IVR held 4.9 mg of ENG and 1.13 mg of EE. *In vitro* release assays for ENG and EE were performed on an orbital shaker table (120 rpm) by placing each type of IVR in 100 mL deionized water at 37°C and daily drug releases were quantified via high-performance liquid chromatography. Use of these assays to define areas under the curve for both these sex steroids indicated N-IVR accumulatively released about 46% less ENG and 39% less EE than NuvaRing® (Fig 1).

**Figure 1.**
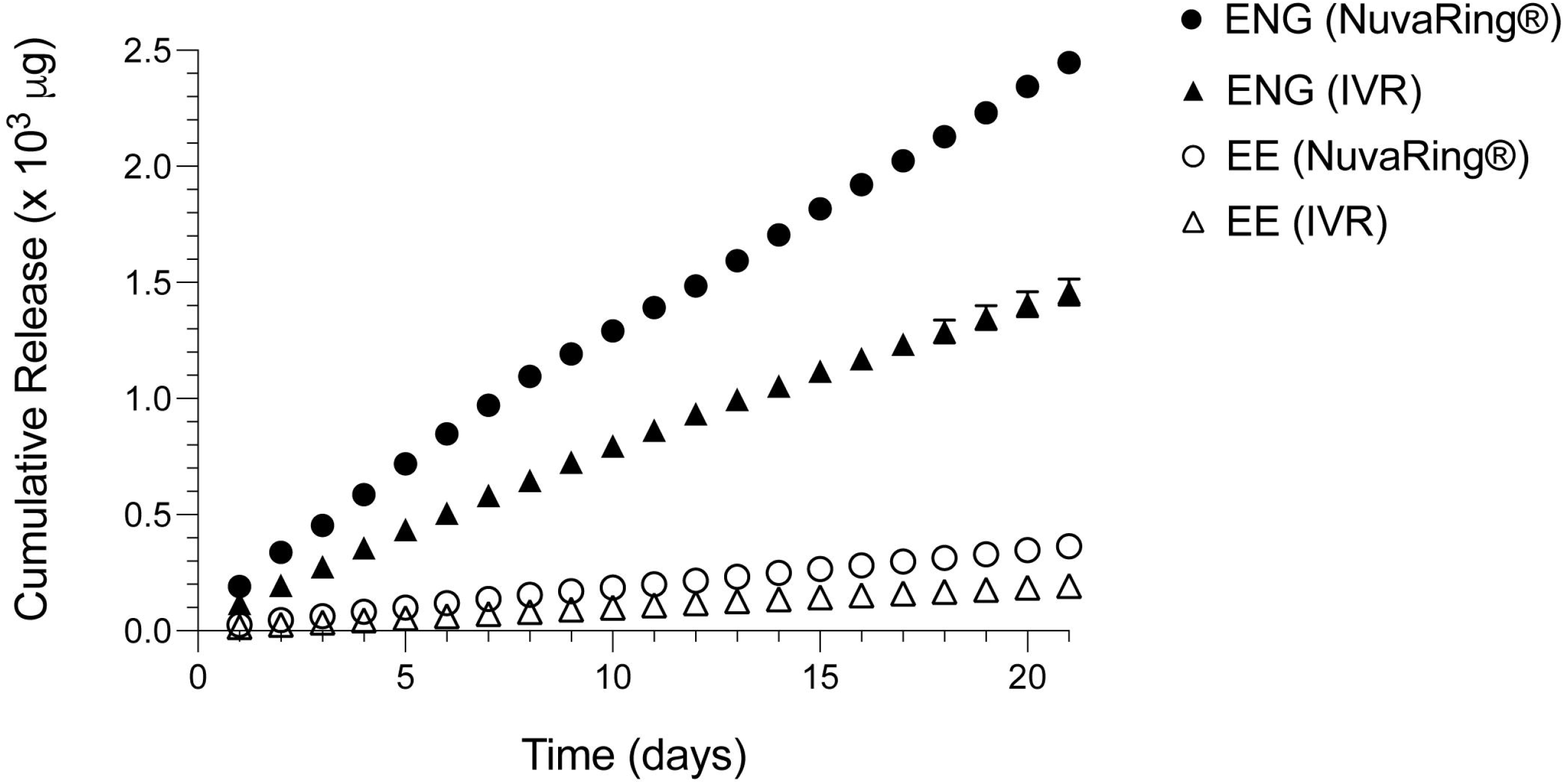
Re-sizing NuvaRings® for use in RM does not impair ENG or EE release. Because N-IVR manufacture involved both cutting and thermal bonding procedures, *in vitro* assays were used to confirm there was sustained release of ENG and EE by N-IVR. In the manufacturing process, NuvaRing® were reduced to 60% of their original size and area under the curve analyses of release assay results showed that the N-IVR released about 46% less ENG and 39% less EE vs. NuvaRing®. ENG, etonogestrel, EE, ethinyl estradiol; N-IVR; NuvaRing® re-sized for RM use; RM, rhesus macaques.

### Animals and procedures

All procedures adhered to American Association for Accreditation of Laboratory Animal Care regulations and were deemed acceptable by the Institutional Animal Use and Care Committee at the University of California, Davis prior to study initiation. At the time this research was performed, all three rhesus macaques (RM) were seven years of age. Observation of these animals in the six months prior to study initiation indicated that each had normal menstrual activity and baseline specimens (i.e., from untreated RM) were collected at variable days after evidence of menstruation was last observed. Peripheral blood and genital biopsy specimens were collected at indicated time points (Fig 2A) after RM had been anesthetized by intramuscular injection of 10 mg/kg of ketamine hydrochloride (Parke-Davis, Morris Plains, NJ) or 0.7 mg/kg of tiletamine hydrochloride and zolazepam (Fort Dodge Animal Health, Fort Dodge, IA). N-IVR were retained in RM for 12 days. Sixteen days after IVR removal, RM were intramuscularly injected with 1.5 mg/kg of DMPA (Pfizer, New York, NY). In the time between N-IVR removal and DMPA injection, RM displayed evidence that menstrual cycle activity had resumed.

### Quantification of serum sex steroids

Endogenous and exogenous sex steroid serum levels were quantified by the Endocrine Technology Support Core at the Oregon National Primate Research Center. A cobas e 411 analyzer ® (Roche Diagnostics, Indianapolis, IN) measured endogenous estrogen (E_2_) and progesterone (P_4_) levels via electrochemiluminescence and a liquid chromatography-tandem triple quadrupole mass spectrometry quantified levels of MPA, EE, and ENG [14]. Quantification limits were 0.020 ng/mL for MPA and ENG, 0.010 ng/mL for EE, and 0.050 ng/mL for E_2_ and P_4_. Of note, clinical reports indicated within 72 hours of NuvaRing® removal, serum EE levels are undetectable[15].

### Genital permeability assays

*Ex vivo* evaluation of genital epithelial barrier function was performed using previously described methods [9, 16, 17]. Briefly, vaginal biopsy tissue was immediately immersed in chilled RPMI-1640 (Corning, Manassas, VA) supplemented with 10% charcoal/dextran-treated fetal bovine serum (R&D systems, Minneapolis, MN). Tissue samples were washed with PBS before transfer to wells in 96-well plates that contained 50 μL of PBS with 62.5 μg of Texas Red labeled dextran (DR) (MW=70,000 Da) and 50 μg of Lucifer yellow CH, lithium salt (LY) (MW=457.2 Da) (both Thermo Fisher Scientific, Eugene, OR). Tissues were incubated for 45 minutes at 37°C in 5% CO_2_ and fixed for 24 hours in 4% methanol-free formaldehyde. After embedding in 6% agarose, 200-300 μm tissue sections were made using a PELCO EasiSlicer™ (Ted Pella Inc., Redding, California) and counterstained with DAPI (4’, 6-diamidino-2-phenylindole, Thermo Fisher Scientific). Sections were examined with a Nikon A1 confocal microscope and LY molecule tissue infiltration (depicted in green channel images included in Fig 4C) was defined via ImageJ software (pixels/10^4^ μm^2^) [18]. To facilitate assessment of LY molecule tissue incursion, white lines were drawn to distinguish vaginal epithelium from underlying lamina propria.

### Histology

Ectocervical biopsies were fixed for 48 hours in 4% formaldehyde (Thermo Fisher Scientific, Rockford, IL) and paraffin-embedded to generate 5 μm sections used for hematoxylin and eosin (H&E) staining. DSG1 protein levels were measured by immunohistochemistry (IHC). As described earlier [16], unstained ectocervical sections were de-paraffinized with xylene and rehydrated in decreasing ethanol concentrations and deionized water. For antigen retrieval, sections were incubated 20 minutes at 95°C in citrate buffer with 0.02% Tween 20 (pH = 6.0). Endogenous peroxidase activity was quenched by 3% hydrogen peroxidase prior to 4-hour incubation at 4°C with 5% normal goat serum (Cell Signaling Technology®, Danvers, MA) and rabbit anti-DSG1 (clone EPR677766(B), Abcam, Cambridge, MA) diluted 1:100 with SignalStain® antibody diluent (Cell Signaling Technology®). Sections were washed, incubated with SignalStain® Boost Detection and SignalStain® BAD Chromogen (both Cell Signaling Technology®), counterstained with hematoxylin, and covered with mounting medium (Cell Signaling Technology®). The H&E-stained and IHC slides were imaged with the NanoZoomer 2.0-RS scanner (Hamamatus Photonics K. K, Hamamatsu City, Shizuoka, Japan) and analyzed via NDP.view2 software (Hamamatsu Photonics K. K) and ImageJ software, respectively. For H&E images, white lines were drawn to highlight the epithelial-stromal interface and delineate keratinized non-nucleated superficial layers from non-keratinized nucleated layers. Epithelial thickness measurements were obtained from suitable sections (at 100 μm intervals). To quantify DSG1 protein levels, areas were randomly selected to convert and display as optical density per 10^4^ μm^2^.

**Figure 2.**
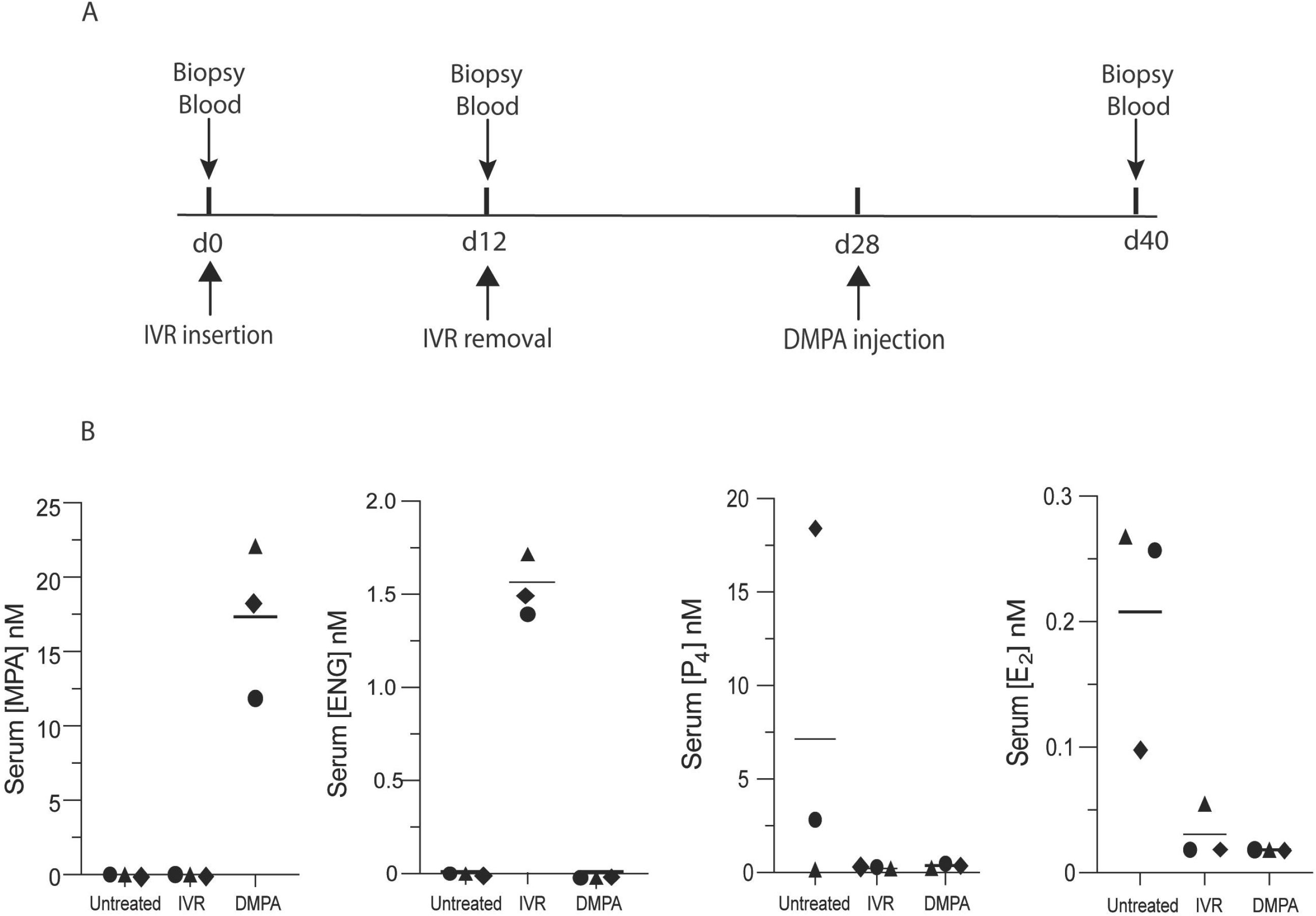
DMPA and N-IVR similarly dampen HPO axis function. A) Timeline for collecting genital biopsy tissue and blood from study animals. B) Left two panels display mean serum levels of MPA and ENG (before and after initiating DMPA or N-IVR); right two panels show comparable mean serum levels of endogenous P_4_ and E_2_ in DMPA- and N-IVR-treated RM. DMPA, depot-medroxyprogesterone acetate; E_2_, estrogen; ENG, etonogestrel; N-IVR, re-sized NuvaRing®; MPA, medroxyprogesterone acetate; P_4_, progesterone; RM, rhesus macaques.

**Figure 3.**
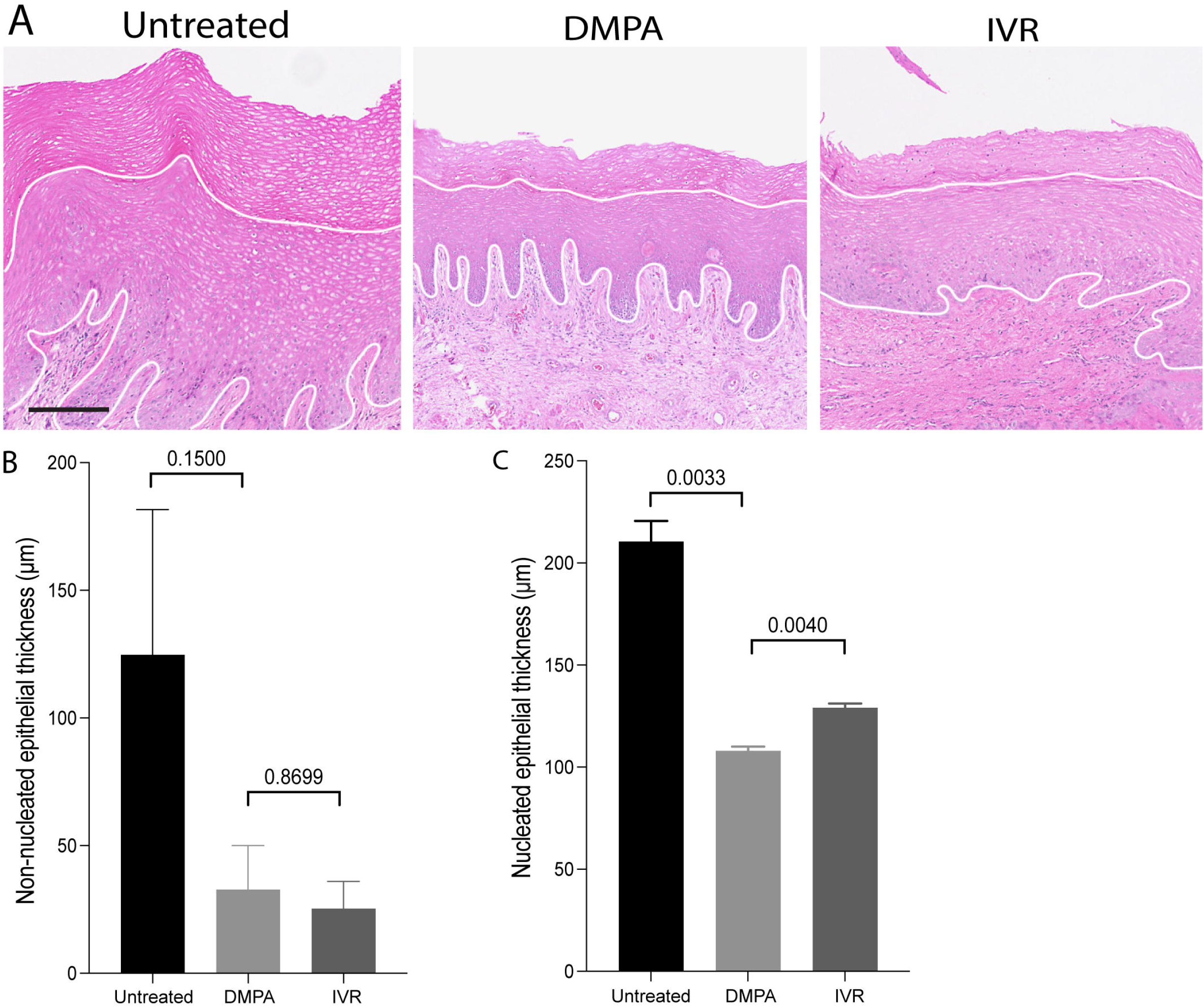
Thinning of nucleated genital epithelium is greater in DMPA- vs. N-IVR-treated RM. A) Representative hematoxylin and eosin (H&E) stained images from genital biopsy tissue before and 12 days after initiating DMPA or N-IVR (scale bar = 100 μm). B-C) Columns compare B) non-nucleated and C) nucleated ectocervical epithelium thickness in RM before and after initiating DMPA or N-IVR. DMPA, depot-medroxyprogesterone acetate; N-IVR, re-sized NuvaRing®; RM, rhesus macaques.

**Figure 4.**
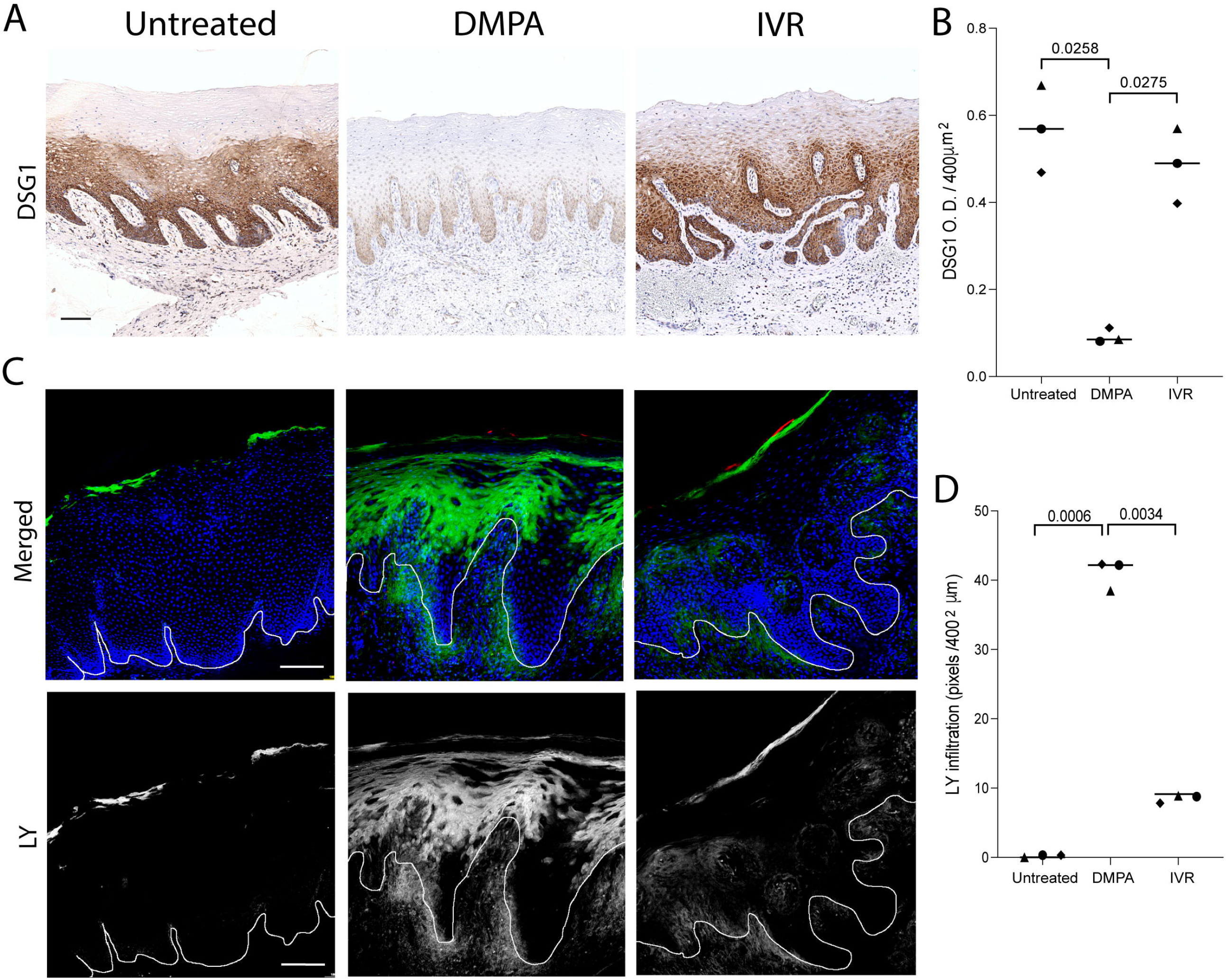
Loss of genital DSG1 protein and increases in genital epithelial permeability are greater in DMPA- vs. N-IVR-treated RM. A) Representative images of immunohistochemical staining for DSG1 protein in vaginal tissue. B) Histograms display the significantly DSG1 protein levels in DMPA- vs. N-IVR-treated RM. C) Representative confocal microscopy images illustrate entry of intravaginally administered LMM molecules into vaginal epithelium. D) Histograms identify significantly greater penetration of LMM molecules into vaginal epithelium of DMPA- vs. N-IVR-treated RM. Scale bars seen in panels A & C = 100 μm; DMPA, depot medroxyprogesterone acetate; DSG1, desmoglein 1; N-IVR, re-sized NuvaRing®; LMM; low molecular mass; RM, rhesus macaques.

### Statistical considerations

All statistical analyses were performed using Prism 9 software (GraphPad, La Jolla, CA). One-way ANOVA and Tukey’s multiple comparisons post hoc test were used for multiple group comparisons. P values ≤ 0.05 were deemed statistically significant.

## RESULTS

### DMPA and N-IVR comparably suppress the HPO axis

The current investigation used 3 reproductive-age RM to compare the effects of DMPA vs. N-IVR on HPO axis function and genital epithelial integrity and permeability. These RM displayed a mean MPA serum level of 17 nM 12 days after intramuscular injection of DMPA (Fig 2B), a value congruent with the 5.8-62 nM MPA range detected in the serum of reproductive age women in the first 2 weeks after intramuscular DMPA injection [19-21]. On the other hand, mean ENG levels in the serum of RM 12 days after N-IVR insertion were approximately 1.5 nM (Fig 2B), a value lower than the 4.0 nM detected 2 weeks after women initiate use of NuvaRing® [15]. Likewise, while serum EE levels measured in women during days 7-21 of NuvaRing® use are about 50 pM [15], we were unable to detect EE in the serum collected from RM 12 days after N-IVR insertion (Fig 2B). Despite serum ENG and EE concentrations lower than those reported in women using NuvaRing®, serum levels of endogenous progesterone (P_4_) and estrogen (E_2_) levels were indistinguishable in RM 12 days after DMPA injection or N-IVR insertion (Fig 2B). The latter results indicated there was comparable HPO axis suppression after systemic DMPA injection or intravaginal (ivag) insertion of the N-IVR.

### Compared to N-IVR, DMPA induces more genital epithelial thinning

Whereas DMPA and N-IVR comparably inhibited HPO axis function (Fig 2B), we posited that combined delivery of progestin and estrogen by N-IVR generates less genital epithelial thinning than intramuscular injection of a progestin alone (i.e., DMPA). Congruent with this hypothesis and prior report [16], non-nucleated and nucleated layers of stratified squamous genital epithelium were significantly thinner in DMPA-treated vs. untreated RM (Fig 3). While mean nucleated epithelium measurements of 207 μm and 132 μm, respectively, in untreated and N-IVR-treated RM suggested that N-IVR likewise promotes genital epithelial thinning, we saw nucleated genital epithelium significantly thinner in RM after DMPA injection vs. N-IVR insertion (Fig 3C). These latter results indicated that unopposed progestins (i.e., DMPA) have greater impact on genital epithelial thickness than combined local release of ENG and EE.

### Compared to N-IVR, DMPA significantly decreases levels of genital DSG1 protein and increases genital epithelial permeability

Because prior studies showed that progestin-mediated loss of genital epithelial integrity is more impactful to barrier function than progestin-mediated loss of genital epithelial thickness [9, 10, 16], we compared levels of the cell-cell adhesion molecule DSG1 in genital tissue from untreated, DMPA-treated, and N-IVR-treated RM. While genital DSG1 protein levels were comparable before treatment and 12 days after N-IVR insertion, we saw significantly lower levels of this desmosomal protein in genital biopsies from DMPA- vs. N-IVR-treated animals (Fig 4A-B). Congruent with the effects on DSG1 protein, permeability of genital epithelium to LMM molecule incursion was comparable before treatment and after N-IVR insertion, whereas DMPA treatment significantly enhanced entry of the LMM molecules (Fig 4C-D). Together, our findings confirm results from previous murine and clinical studies that showed DMPA promotes loss of genital epithelial integrity and barrier function [9, 11, 12], and newly identify that DMPA is more likely than an IVR releasing ENG and EE to impair this first-line host defense in the female genital tract.

## DISCUSSION

Regions of SSA more profoundly impacted by HIV are dually characterized by high rates of HIV infection in AGYW and heavy reliance of this cohort on DMPA for contraception [22, 23]. However, no causal connection between DMPA and HIV susceptibility is established and the extent to which this injectable progestin impacts the high rate of HIV infection in AGYW in SSA is uncertain. Interestingly, injectable contraceptives were first identified as a significant HIV risk factor more than three decades ago [24] and systematic review and meta-analysis of eligible observational studies estimated the risk of HIV is 40% higher in women using DMPA vs. no hormonal contraception [5]. Likewise, while the ECHO Trial found no statistically significant increased rate of HIV infection among women randomized to DMPA, LNG-releasing subdermal implant, or copper intrauterine device, it saw HIV acquisition roughly 30% higher among women using DMPA vs. LNG implant (P = 0.06) [8]. Moreover, this inability to detect statistically significantly (i.e., P ≤ 0.05) increased rates of HIV infection in study participants randomized to DMPA must be evaluated in context of a trial that was powered to detect significance between-group differences if a particular contraceptive increased HIV acquisition by at least 50% [8, 25].

In addition to the substantial financial resources needed to conduct appropriately powered clinical trials, other complexities associated with using clinical research to define the effects of hormonal contraception on HIV susceptibility include ethical constraints that preclude randomizing women to use no form of long-acting contraceptive and the inability to fully eliminate the residual confounding created by increased frequencies of unprotected sex in women using hormonal contraception vs. no contraception. These obstacles are not trivial and highlight the continued reliance on experimental models to both define the impact of exogenous sex steroids on mechanisms of antiviral defense and identify contraceptive choices least likely to enhance HIV transmission [9]. The current study provides clear illustration of the utility of animal models, identifying intramuscular DMPA injection of RM significantly weakens genital epithelial integrity and barrier function and that N-IVR is less likely to induce these changes. These findings also imply that IVR releasing ENG and EE may be less likely than DMPA to promote sexual transmission of HIV and other genital tract pathogens, and this possibility is an area of active research in our laboratory.

Though RM are a relevant preclinical model for defining effects of exogenous sex steroids in the female genital tract, there are limitations associated with our research. Primarily, though comparing the effects of DMPA and EEE-IVR on genital epithelial integrity and barrier function in the same RM is a strength of our study design, conclusions drawn from these results may be limited by the small number of RM examined. As another potential limitation, serum ENG and EE concentrations in N-IVR-treated RM were less than those seen in women at similar timepoints after initiating NuvaRing® use. Specifically, while 4.6 nM of ENG and 67 pM of EE are the approximate values measured in the serum of women 2 weeks after NuvaRing® insertion [15], RM in the current investigation displayed a mean ENG level of 1.6 nM and undetectable EE levels 12 days after N-IVR insertion. Interestingly, prior studies indicate that MPA is degraded more quickly in macaques than humans, displaying terminal elimination half-lives of roughly 9 vs. 50 days respectively [19, 26-29]. As NuvaRings® were reduced to 60% of original size and RM are roughly 20% the weight of a typical adult women, our data also imply there is more rapid degradation of ENG and EE in RM vs. humans. Despite the lower than anticipated serum levels of ENG and EE in our investigation, circulating levels of endogenous E_2_ and P_4_ were comparably reduced by DMPA and N-IVR. These findings indicate DMPA and N-IVR comparably suppress HPO axis function and that comparison of the effects of these treatments on genital epithelial integrity and barrier function in RM is meaningful.

Current findings thus provide fresh evidence that unopposed progestin (e.g. DMPA) is more likely than combined release of progestin and estrogen to weaken genital epithelial integrity and barrier function, key host defense mechanisms in the female genital tract that reduce the risk of virus acquisition.

## ACKNOWLEDGEMENTS

This research was supported by the Eunice Kennedy Shriver National Institute of Child Health and Human Development (Award Number R01HD094634) and CNPRC base grant P51OD011107. The authors are solely responsible for the content of this publication, which does not necessarily represent official views of the National Institutes of Health. Quantification of endogenous and exogenous sex steroids levels in serum samples was performed by the Endocrine Technology Support Core at the Oregon National Primate Research Center (Award Number P510D011092). Authors are grateful for CNPRC staff that provided excellent animal care in the midst of the COVID-19 pandemic.

